# Interpretable machine learning meets systems biology to decode genotype-phenotype maps

**DOI:** 10.64898/2026.03.16.712082

**Authors:** R.M. Rajeeva Lokshanan, Roshan Balaji, Himanshu Sinha, Nirav Pravinbhai Bhatt

## Abstract

Resolving causal genes from quantitative trait loci (QTL) remains fundamentally limited by linkage disequilibrium. We developed an interpretable machine learning framework that captures higher-order nonlinear genotype-phenotype relationships and allows conditional evaluation of genetic variants, enabling statistical decorrelation of linked loci. Applied to *Saccharomyces cerevisiae* segregants across chemical stress conditions, our method achieved >75% prediction accuracy and identified known causal genes, including *MKT1* (genotoxic stress) and *IRA2* (osmotic stress). SHAP-based analysis recovered 56% of the validated pleiotropic genes, compared with 36% by conventional contingency testing. Integration with genome-scale metabolic models revealed pathway enrichments distinguishing high-growing strains, including carbon transport, glycolysis, and oxidative phosphorylation. Notably, gene regulatory network analysis identified a novel function for *PDR8* in protein mannosylation and cell wall integrity—functions extending beyond its role in drug resistance. This framework demonstrates that interpretable machine learning, coupled with systems biology, transforms QTL associations into mechanistic biological insight.

## INTRODUCTION

A central objective of modern genetics is to elucidate the complex relationship between an organism’s genotype and its resulting phenotype. Widely used approaches toward this goal include Quantitative Trait Loci (QTL) mapping in model organisms^1–3^ and Genome-Wide Association Studies (GWAS) in human populations^4,5^. In practice, both frameworks rely predominantly on additive, marginal association models to link sequence variants to phenotypic variation. While these models have proven effective for identifying loci associated with traits, they face fundamental limitations in resolving causal variants with high precision. A principal challenge arises from linkage disequilibrium (LD), the non-random association of alleles at different genomic loci. Variants in LD are inherited together, resulting in highly correlated genotypes across linked loci. Consequently, multiple variants within the same region often yield indistinguishable statistical signals, making it difficult to disentangle the true causal variant from nearby variants that merely co-segregate with it ^6^. This lack of identifiability under LD represents an intrinsic limitation of marginal association testing and complicates downstream biological interpretation.

Several methodological strategies have been proposed to address these challenges. Fine-mapping approaches perform post hoc analyses of associated regions to prioritize putative causal variants. These methods, such as SuSiE^11^, FINEMAP^12^, and CAVIAR^13^, formulate the identification of causal variants as a ‘variable selection’ problem, wherein the goal is to identify variants or sets of variants that are likely to be causal for a given phenotype or trait. While effective at narrowing to probable causal genes, these methods are typically applied per-phenotype and rely on additive modeling assumptions, limiting their ability to identify variants that exert pleiotropic effects across multiple traits or participate in non-additive genetic interactions. As a result, epistatic and context-dependent effects may remain undetected. An alternative class of methods employs machine learning models trained on biological and functional annotations to rank variants within associated regions^14–17^. Although these approaches can incorporate richer feature representations, their performance depends critically on the availability of large, high-quality curated training datasets, which are not uniformly available across tissues, populations, or phenotypes. This dependence introduces annotation bias and limits their generalizability. A third line of work seeks to improve causal interpretation by integrating multi-omic datasets, including transcriptomic, epigenomic, and chromatin interaction data, to better contextualize loci identified by QTL and GWAS analyses^6,9,18,19^. While conceptually powerful, such integrative approaches are constrained by the limited availability, heterogeneity, and variable quality of matched multi-omic datasets, posing a substantial barrier to their broad application.

In this work, we use an interpretable machine learning approach that overcomes LD-induced confounding by exploiting the non-linear, multi-locus architecture of genotype-phenotype relationships. Genes do not act in isolation, and our model utilizes the fact that interacting genes may not be genetically linked to one another. Applied to *Saccharomyces cerevisiae* chemical stress phenotypes^1,20,21^, our method resolves quantitative trait genes (QTGs) with high precision, identifies pleiotropic hub genes, and integrates with constraint-based metabolic and regulatory network models to provide mechanistic insight. This combined framework revealed a previously unknown role for the transcription factor *PDR8* in protein mannosylation and cell-wall integrity, extending its function beyond drug resistance. Our results demonstrate that interpretable ML, when coupled with systems biology approaches, can transform statistical associations into a mechanistic understanding of genotype-phenotype relationships.

## RESULTS

### Nonlinear genotype-phenotype modeling identifies causal genes within linked QTL regions

We hypothesized that a model capable of accurately predicting phenotype from genotype would facilitate the identification of causal genes through interrogation of its decision-making mechanisms. To test this, we developed a unified model that integrates genotype information with chemical latent representations to predict phenotypes across diverse chemical environments (Figure 1B).

**Figure 1:**
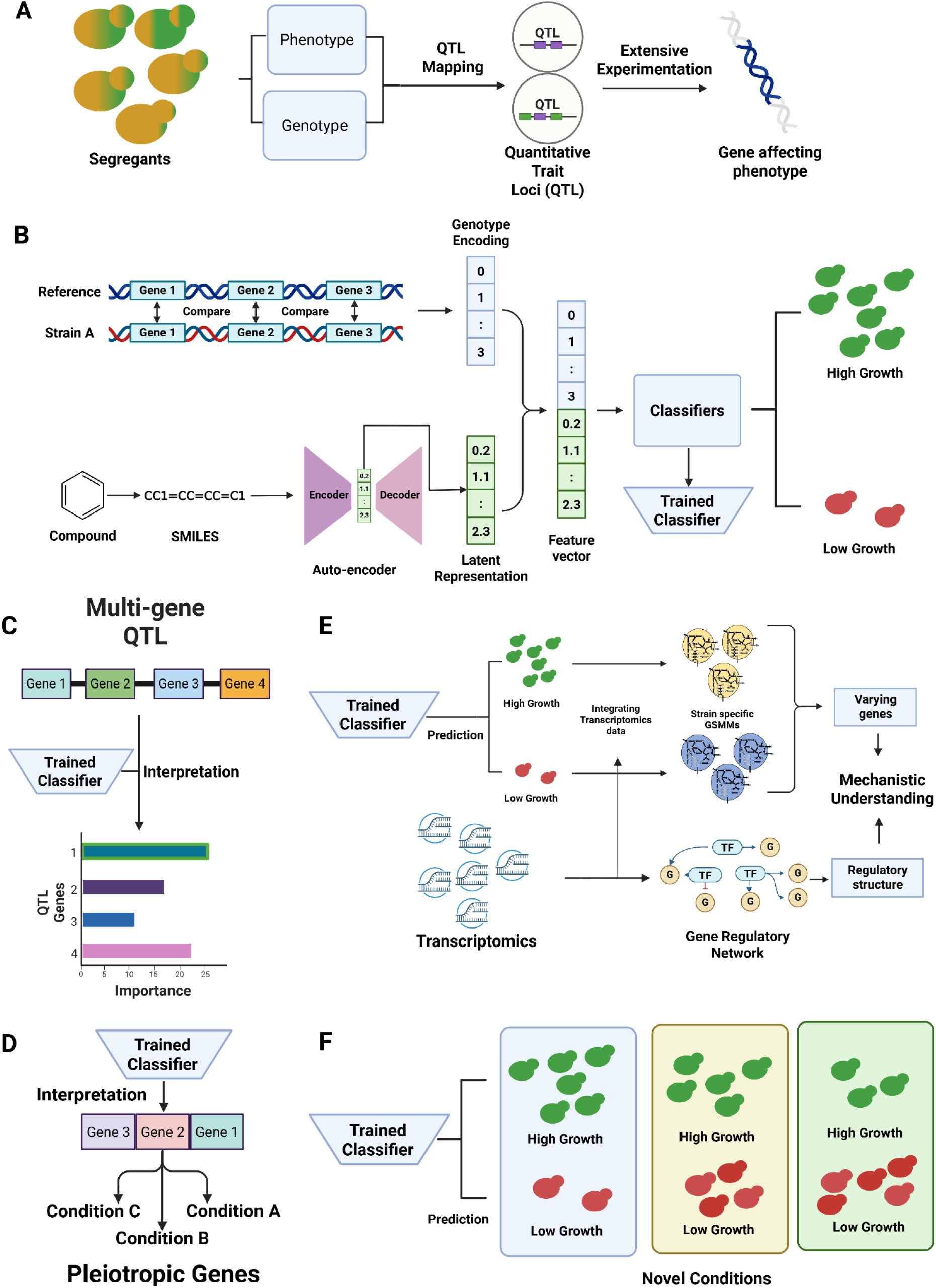
Interpretable ML framework for QTG identification. (A) The ML framework for jointly modelling the genotype and chemical environment information. (B) Schematic comparing standard QTL mapping (yielding broad intervals) with our GBDT+SHAP approach for resolving causal genes. (C) Detection of pleiotropic genes affecting multiple phenotypes. (D) Integration with constraint-based models (GSMM, GRN) for mechanistic interpretation. (E) Framework for prediction in novel chemical environments.

To study genotype–phenotype relationships and their interactions with environmental conditions, we used yeast segregant population data derived from crosses between the BY and RM strains from three studies (hereafter referred to as Bloom2013, Bloom2015, and Bloom2019)^20–22^. These studies include both chemical perturbations and abiotic conditions such as pH and temperature; here, we restricted the analysis to chemical environments. The phenotype is the growth of the segregants in the specific chemical environment.

The raw dataset contains thousands of genetically distinct segregants phenotyped across 50 chemical conditions (Supplementary Figure S1A). Approximately 30,000 variants differ between the BY and RM strains. We restricted our analysis to coding variants, as the functional consequences of variants in non-coding regions are more difficult to predict, and phenotypic effects are primarily mediated through proteins. After preprocessing, the genotype data consisted of SNPs affecting approximately 3,000 genes.

In total, the dataset includes 50 unique chemicals that primarily differ in growth medium composition. To incorporate environmental information into the machine learning models, we generated a 256-dimensional latent representation for each chemical from its SMILES string using a deep-learning autoencoder (Supplementary Figure S1B). The model was adopted from a previous study and pretrained on around 114 million compounds from the PubChem database to learn compact molecular embeddings^23^. This representation allows the model to learn common genetic effects across chemically related environments.

Genotype features were defined based on genes containing coding variants, while chemical features were derived from the learned SMILES embeddings. Integrating these representations enabled the model to simultaneously capture environment-specific genetic effects and cross-environment genetic drivers, facilitating the identification of quantitative trait genes (QTGs) (Figure 1C–D).

Applying our model to the datasets, we observed that gradient-boosted decision tree (GBDT) models achieved mean cross-validation AUC-ROC exceeding 75% across chemical stress conditions, substantially outperforming random baselines and alternative machine learning models (Supplementary Figure S3A). Unlike traditional methods such as marginal association tests^6,9^, GBDT models capture linear and higher-order nonlinear genotype-phenotype relationships with each genetic variant conditioned on all others, enabling statistical decorrelation of linked loci. To interpret model predictions and prioritize candidate causal genes within mapped QTL intervals, we applied SHAP (SHapley Additive exPlanations) analysis to quantify the contribution of each gene to phenotype prediction for each chemical condition (Figure 1C).

We validated our framework across multiple chemical conditions to infer environment-specific effects, identifying QTGs within each QTL (Figure 2). Under 4-nitroquinoline-N-oxide (4NQO) genotoxic stress, SHAP analysis identified *MKT1* (YNL085W) as the primary causal gene within a locus on chromosome XIV (Figure 2A). This locus explains approximately 15% of phenotypic variance but was previously unresolvable due to high LD^20^. Notably, the causal role of *MKT1* has been independently validated through allele replacement experiments^24^. Within the same condition, SHAP identified *MLH2* (*YLR035C*), in a locus in chromosome XII, which encodes a DNA mismatch repair component—mechanistically consistent with the established DNA-damaging activity of 4NQO^25^ (Figure 2B). For osmotic stress (sorbitol), we resolved *IRA2* (*YOL081W*) as the principal causal gene within a chromosome XV locus (Figure 2F). *IRA2* encodes a GTPase-activating protein that negatively regulates the RAS-cAMP signaling cascade, a central coordinator of yeast osmotic stress responses^26^. At another locus on the same chromosome (Figure 2G), we extracted *VHS3* (*YOR054C*), a non-essential gene shown to confer reduced resistance to NaCl, another osmotic stressor^27^. In a chromosome IV locus (Figure 2E), the top-ranked gene was *FAR11 (YNL127W)*. FAR11 is involved in the recovery of cells from cell cycle arrest and has been associated with stress response^28^, although its role in osmotic stress remains undocumented. This locus also contains other genes which have been implicated in other stresses, such as *KRE33* (YNL132W)^29,30^ and *NRK1* (YNL129W)^31,32^, but not in the context of osmotic stress. Congo red stress, which disrupts cell wall assembly, identified two primary loci (Figure 2C–D). On chromosome XIII, SHAP identified *BUL2* (*YML111W*), whose paralog *BUL1* (YMR275C) is a known cell wall regulator^33^, confirming the method’s ability to recover established biology. Interestingly, on chromosome II, *DSF2* (*YBR007C*), a gene of poorly characterized function, emerged as the top candidate, suggesting a novel role in cell wall biology. For neomycin resistance, *CDC5* (*YMR001C*), a polo-like kinase essential for cell cycle progression, was identified as the primary candidate (Figure 2H). *CDC5* has documented sensitivity to bleomycin^34^ but not previously to neomycin, suggesting broader antibiotic response functions. A detailed description of all the previously identified QTLs from the Bloom2013 dataset and the resolved QTGs is available in Supplementary File 1.

**Figure 2:**
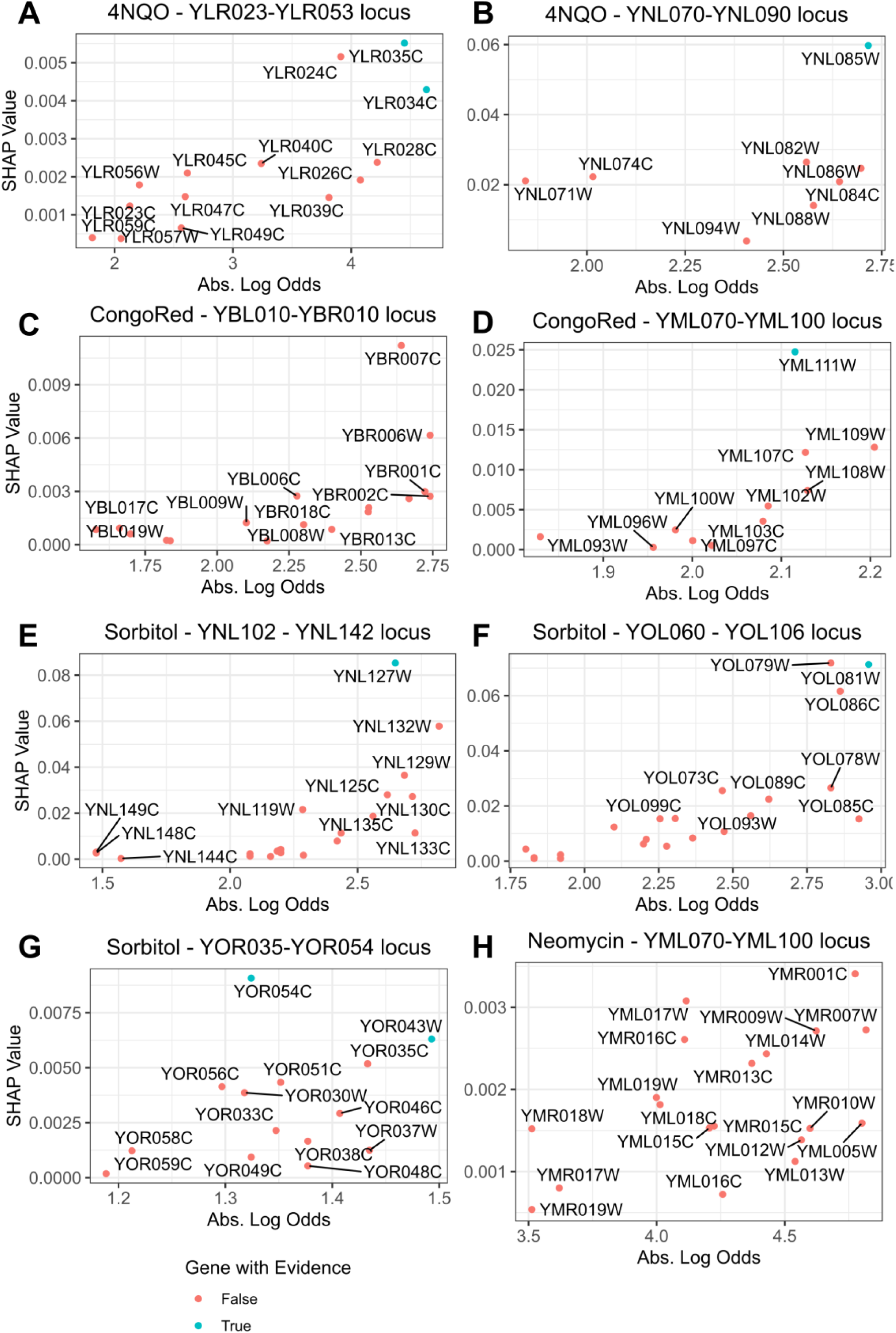
Validation of causal gene identification across chemical stress conditions. SHAP values (y-axis) versus absolute log odds ratio from contingency analysis (x-axis) for genes within QTL intervals. (A–B) 4NQO genotoxic stress: *MKT1* and *MLH2* identified as top candidates. (C–D) Congo red cell wall stress: identification of *DSF2* and *BUL2*. (E–G) Sorbitol osmotic stress: *IRA2* resolution. (H) Neomycin: *CDC5* identification.

### SHAP analysis improves recovery of pleiotropic genes

The adaptability of organisms to multiple chemical conditions suggests mechanisms that allow organisms to modify their biological pathways in response to a changing environment. Rather than relying on a unique set of genes in each environment, organisms have evolved such that a subset of genes is shared across multiple conditions^35^. Genes that affect multiple phenotypes are termed pleiotropic genes. Pleiotropic genes are important because they often act as hub genes that control multiple aspects of the organism^36,37^. Analysing them can improve our understanding of the genetic architecture underlying phenotypic variation. However, pleiotropic genes are difficult to identify using marginal tests because linkage disequilibrium (LD) confounds variants across conditions. Training our model across all chemical conditions simultaneously enabled the identification of features important across multiple stress responses (Figure 1D).

To ensure a valid comparison, we performed a contingency analysis across all chemical conditions in a given dataset rather than per chemical condition. In the Bloom 2013 dataset^20^, genes relevant across multiple conditions have higher SHAP values, indicating that the model can identify pleiotropic genes (Figure 3A). This indicated that the model was using pleiotropic genes for predicting the phenotype. To obtain a more confident evaluation of how well the model identified pleiotropic genes, we identified the causal pleiotropic genes from the Bloom2019^1^ data. Comparing SHAP-based identification to contingency analysis revealed substantially improved recovery of known pleiotropic effectors. SHAP identified 56% (35/63) of previously validated pleiotropic genes (Figure 3C), compared to 32% (20/63) by Fisher’s exact test with Benjamini-Hochberg correction (FDR <0.05) (Figure 3B). Notably, SHAP recovered genes missed by contingency analysis, including *SAL1 (YOL083W)*, an ADP/ATP transporter, and *KRE33 (YNL132),* an acyltransferase required for the biogenesis of the small subunit component of the ribosome, both of which ranked among the top 10 genes by aggregate SHAP value across conditions.

**Figure 3:**
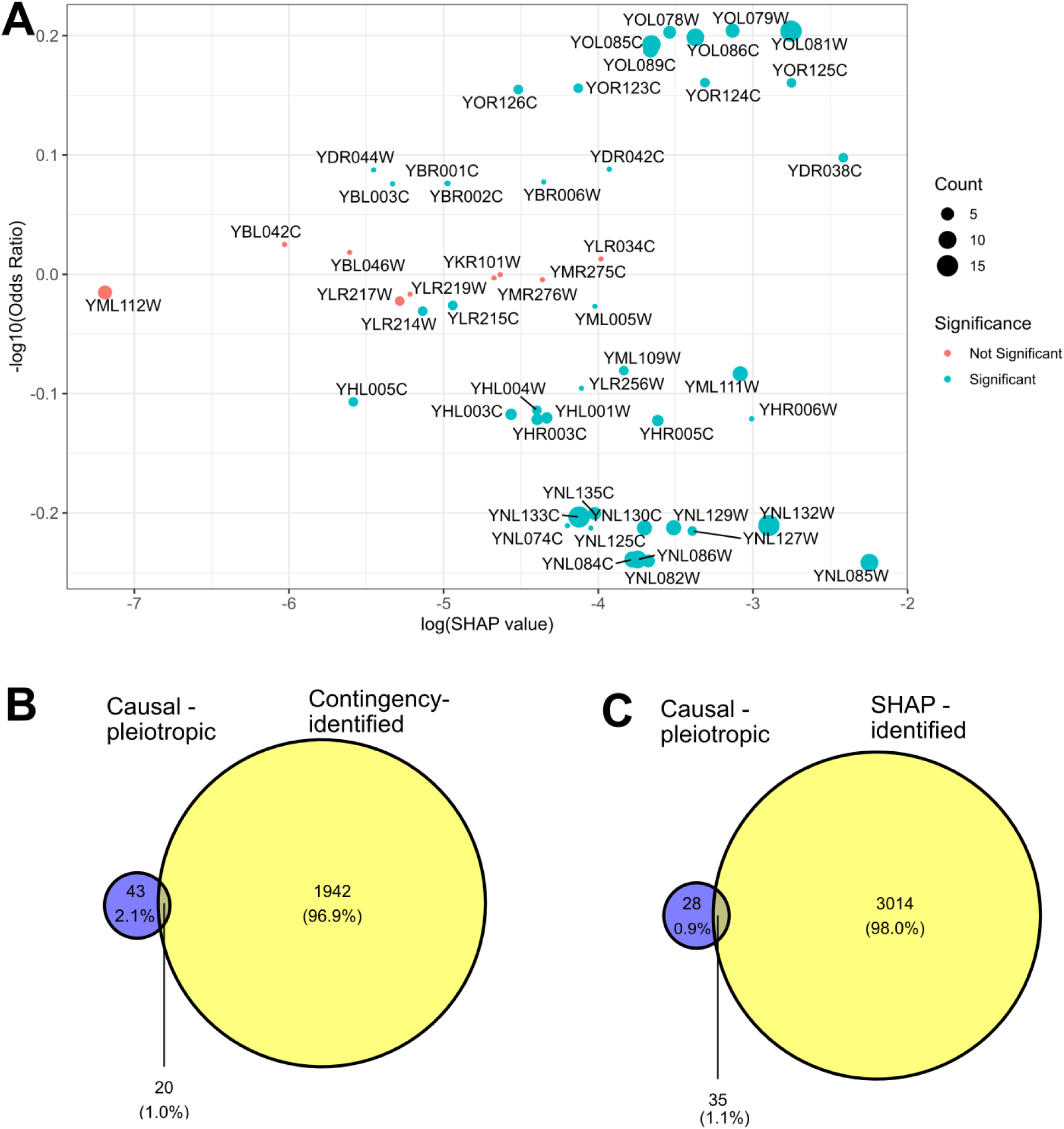
Superior recovery of pleiotropic effector genes. (A) Relationship between SHAP values and contingency table odds ratios for pleiotropic genes across conditions. Point size indicates the number of conditions implicated; color indicates Fisher’s exact test significance (FDR < 0.05). (B–C) Venn diagrams showing pleiotropic gene recovery in Bloom2019 data: SHAP recovers 56% (35/63) vs. contingency analysis 36% (20/63).

### Metabolic flux analysis identifies pathways underlying growth variation

To complement statistical association and enable mechanistic understanding, we constructed strain-specific genome-scale metabolic models (GSMMs) using the yeast-GEM consensus model^38^ (Figure 1E) and transcriptomic data from the Bloom2013 dataset grown in YNB media^39^. Model-predicted growth correlated significantly with experimental fitness (Spearman ρ = 0.31, P <10⁻^16^) (Figure 4A), validating the metabolic reconstructions. Parsimonious flux balance analysis (pFBA), which analyzes metabolic models to identify genes that are active and required for maximal growth^40^, identified 194 genes consistently active for growth in high-growing strains versus 44 genes in low-growing strains, of which 42 were also active in high-growing strains. Unsurprisingly, the 152 genes unique to high-growing strains were enriched for carbon transport (28.3-fold enrichment, P <0.05), glycolysis (17.6-fold enrichment, P <0.05), oxidative phosphorylation (11.9-fold enrichment P <0.05), and nucleotide biosynthesis (7.2-fold enrichment, P <0.05), the processes known to contribute to cellular growth (Figure 4C, Supplementary File 2). To investigate whether the increased glucose uptake leads to changes in downstream reactions in yeast, a reaction-level analysis was performed using a flux-sampling approach^41^. The analysis identified 82 reactions uniquely active in high-growing strains, enriched for amino acid biosynthesis pathways including phenylalanine, tyrosine, and tryptophan metabolism (>20-fold enrichment, P < 0.05) (Figure 4B). Taken together, the flux patterns indicated that high-growing strains couple increased carbon transport with elevated flux through central metabolism, resulting in greater biomass production.

**Figure 4:**
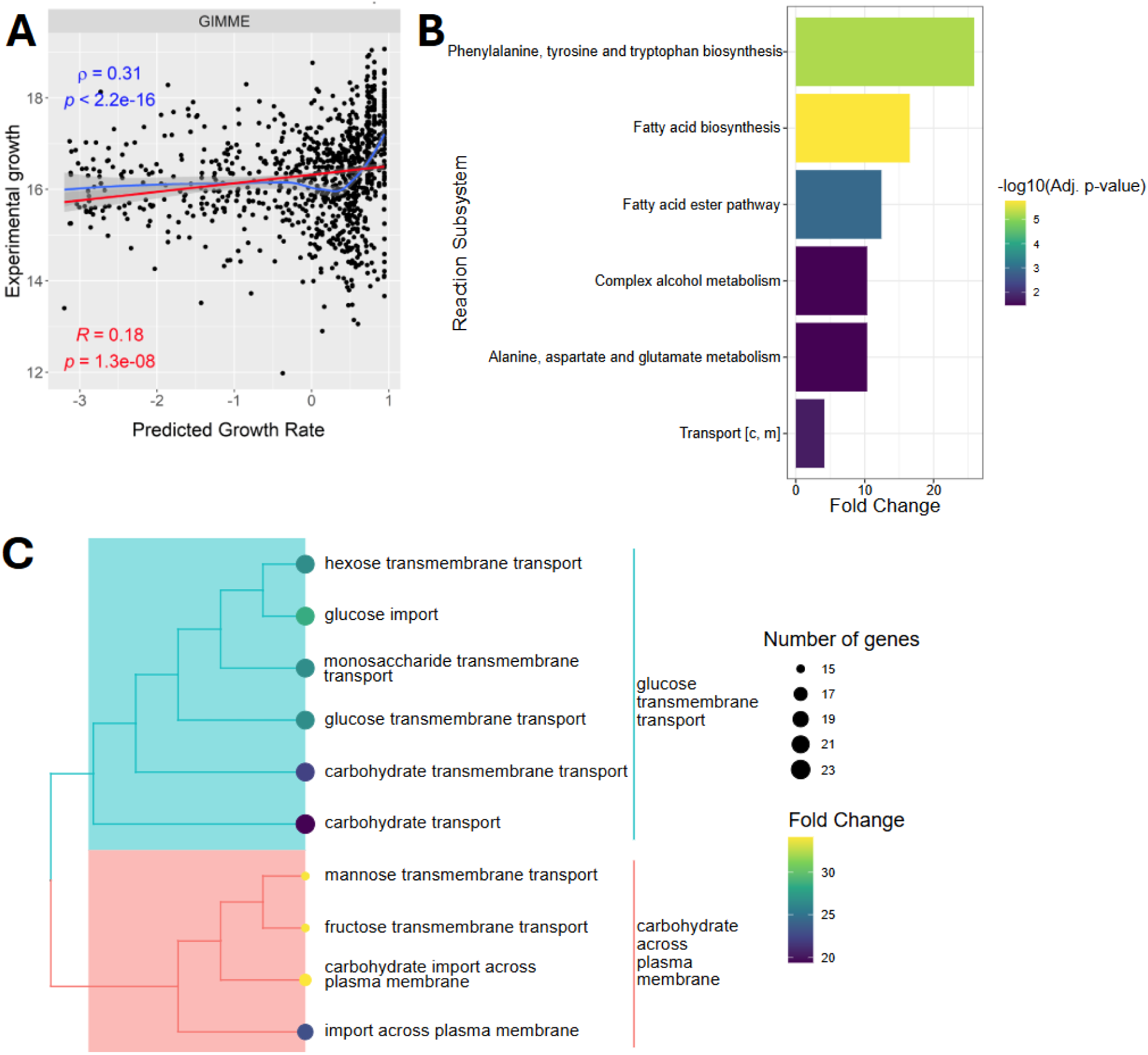
Genome-scale metabolic model analysis identifies growth-associated pathways. (A) Validation of strain-specific GSMMs: model-predicted growth versus experimental fitness (Spearman ρ = 0.31, P < 0.05). (B) Subsystem enrichment for reactions unique to high-growing strains. (C) Top 10 GO pathway enrichments for genes uniquely active in high-growing strains.

### Gene regulatory network analysis reveals novel *PDR8* function in protein mannosylation

Gene regulatory networks (GRNs) provide a complementary framework for representing regulatory relationships between regulatory signals and metabolic outcomes. Incorporating GRNs in this workflow enables integrating insights from SHAP-based explainability and contingency table analyses with findings from GSM modeling. To facilitate this integration, a unified GRN was constructed using the same gene expression data used to build the strain-specific models^39^ and curated yeast regulatory proteins from YEASTRACT^42^. Network inference was performed using the BioNERO package^43^ for Bloom2013 strains grown in YNB medium. The resulting GRN had 245 transcription factors, 4,919 targets, and 60,991 edges. Seventy common genes were identified by SHAP-based explainability analysis, and the transcription factors in the constructed GRN. We focused on a subnetwork extracted from the constructed GRN that met the following criteria: its nodes consisted of the common transcription factors identified by the SHAP analysis and the targets identified by pFBA analysis in high-growing strains. The regulation of gene expression in response to nitrogen and carbon stimuli was among the most highly enriched pathways among the transcription factors in this subnetwork (Figure 5A). The intersection of these orthogonal approaches substantially increased confidence in regulatory predictions. Notably, *YLR266C* (*PDR8*), a transcription factor previously characterized mainly for drug resistance^44^, showed differential regulation of five genes (Figure 5B): *PMT1* (*YDL095W*), *PMT3* (*YOR321W*), *PMT5* (*YDL093W*), *ADE16* (*YLR028C*), and *NMA2* (*YGR010W*), which were highly enriched for protein mannosylation pathways (Enrichment Fold Change> 200, P <0.05). This was because three of these, *PMT1*, *PMT3*, and *PMT5*, encode O-mannosyltransferases essential for cell wall integrity and protein quality control in the endoplasmic reticulum^45,46^. This finding suggested that *PDR8* confers chemical resistance through cell wall maintenance rather than solely through direct regulation of drug efflux transporters, a function previously unlinked to this transcription factor.

**Figure 5:**
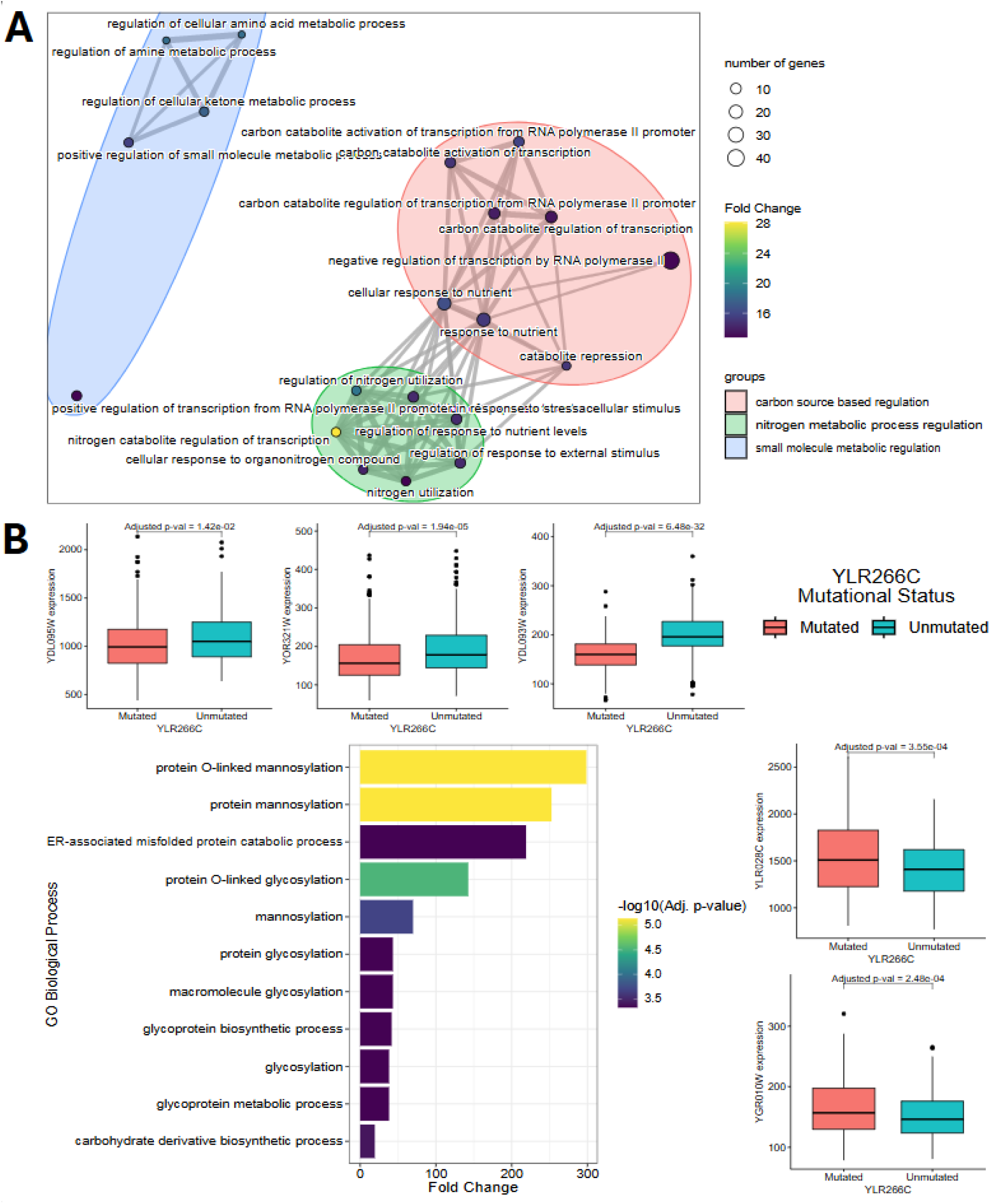
Gene regulatory network reveals novel *PDR8* function. (A) Enriched GO terms among GRN regulators intersecting with SHAP-identified genes. Edge thickness indicates semantic similarity between terms. (B) *PDR8* regulatory network showing downstream targets *PMT1*, *PMT3*, *PMT5*, *ADE16*, and *NMA2*, implicating *PDR8* in protein mannosylation.

### Learned chemical representations enable prediction in novel environments

Finally, we studied the generalization capacity of our framework across datasets and chemical conditions (Figure 1F). A single model trained on the Bloom2015 dataset (18 chemical conditions) and tested on the Bloom2013 dataset (39 chemical conditions) demonstrated the capability of the model to learn from one dataset and predict on another. Models combining both chemical and genotype data outperformed genotype-only models in predicting response to chemicals absent from training data (Figure 6A; Supplementary Figure S5). At the chemical level, information learned from cobalt chloride and magnesium chloride supported accurate predictions for calcium chloride (AUC-ROC = 0.61) and magnesium sulfate (AUC-ROC = 0.59), respectively. Likewise, patterns learned from xylose, raffinose, and trehalose enabled the prediction of responses to sorbitol (AUC-ROC = 0.73) and galactose (AUC-ROC = 0.63). Raffinose and xylose performance remained high as expected as they are present in both the train and test datasets. UMAP visualization of chemical embeddings revealed clustering by chemical similarity, explaining this transfer capacity (Figure 6B). We also verified that this observation was not limited to this particular combination of train and test datasets and observed that the transferability applied to other combinations too (Supplementary Figures S3B, S4-S8). These results demonstrate that our framework can generate testable hypotheses for organism responses in data-limited scenarios and previously uncharacterized chemical environments.

**Figure 6:**
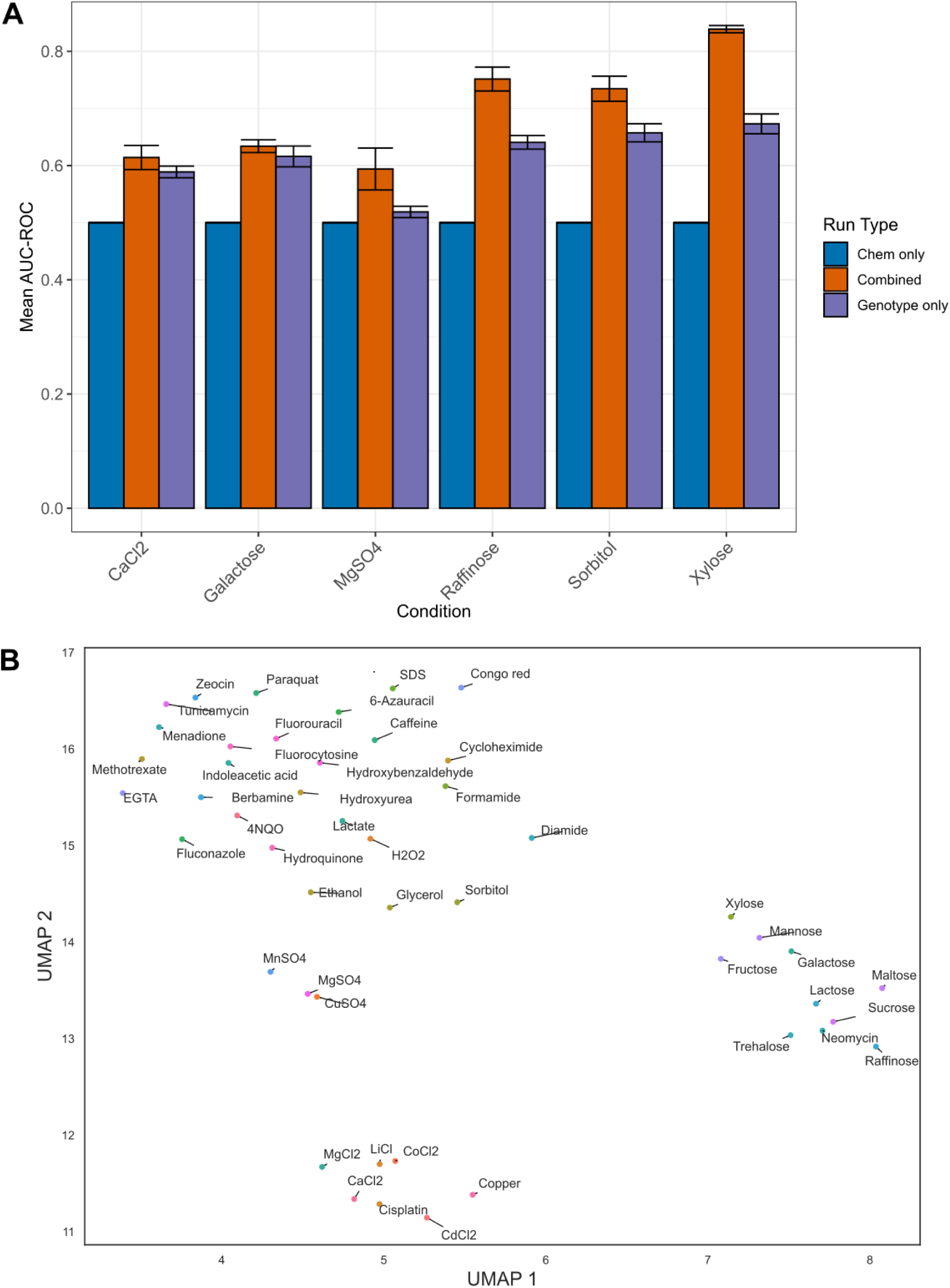
Framework generalization across chemical conditions. (A) Selected condition-wise prediction accuracy for models trained on Bloom2015 (18 conditions) and tested on Bloom2013 (39 conditions), demonstrating transfer to held-out chemicals. The full figure is Supplementary Figure S5. (B) UMAP projection of chemical embeddings showing clustering by chemical class, explaining cross-condition generalization.

## DISCUSSION

Our integrative framework demonstrates that interpretable machine learning can overcome a long-standing limitation of quantitative genetics: distinguishing causal variants from linked markers within QTL intervals. By leveraging the ensemble structure of gradient-boosted decision trees—where each feature is evaluated conditionally on all others—our approach enables statistical decorrelation of linked loci, a capability that marginal association tests lack. Previous studies have applied machine learning to rank causal genes within QTLs^14,16,17^; however, these approaches are typically trained on curated sets of known QTGs and organism-specific genetic features, most notably in well-studied systems such as *Arabidopsis thaliana* and *Sorghum bicolor*. In contrast, our model is trained solely to predict phenotype from genotype, with QTG identification emerging during post hoc model interpretation. This design reduces organism-specific assumptions and makes the framework applicable to less-studied organisms where prior knowledge of causal genes is limited or unavailable.

The validation of known QTGs (*MKT1* in 4NQO, *IRA2* in sorbitol) alongside mechanistically relevant novel candidates (*MLH2* in DNA damage, *BUL2* in cell wall stress) suggests that SHAP-based interpretability provides a biologically meaningful signal rather than a statistical artifact. Pleiotropy detection is notoriously difficult with marginal tests because of LD confounding compounds across phenotypes, and the superior pleiotropic effector recovery achieved with the GBDT framework (56% versus 32%) has practical implications for prioritizing experimental targets. The identification of *IRA2* as a regulator of osmotic stress and a pleiotropic gene affecting growth across multiple conditions, and its role as a RAS-cAMP negative regulator, directly connects our prediction to established yeast stress response circuitry.

The novel role of *PDR8* in protein mannosylation exemplifies the power of integrating statistical association with mechanistic modeling. Previously characterized solely for drug resistance, *PDR8* mutants exhibit dysregulation of *PMT1/3/5*, suggesting broader regulatory function in cell wall integrity. This generates the testable hypothesis that *PDR8* confers chemical resistance through cell wall maintenance rather than by directly regulating transporters.

The contingency table analysis used to identify QTLs compares the presence or absence of a mutation within a gene and a binary phenotype. This means if a gene has no mutation, we will not be able to discern its function even if it is pertinent to the phenotype. This is in contrast to interval mapping, which can implicate genomic regions even when the causative genes themselves are unmutated by leveraging linkage to neighboring variants associated with the phenotype^47^. While our method mitigates LD confounding, it cannot fully eliminate it; variants in perfect LD remain indistinguishable without experimental perturbation. The binarized phenotype requirement may sacrifice information in continuous traits. Performance depends on adequate sample size; the 1,012 segregants represent an unusually large population.

Extension to human GWAS faces additional challenges: larger genomes, smaller effects^48^, and complex population structure^49^; yet the fundamental principle should still hold. Integration with eQTL data and protein interaction networks could further enhance resolution. In summary, our approach combines ‘top-down’ machine learning modeling with ‘bottom-up’ systems biology. It demonstrates that the genotype-phenotype map can be investigated with greater resolution and mechanistic insight than traditional approaches currently allow.

## METHODS

The proposed framework integrates quantitative trait locus (QTL) mapping with interpretable machine learning to mitigate LD and improve understanding of GP relationships (as shown in Figure 1). The workflow begins with the segregant population and their phenotype data, followed by traditional QTL mapping to identify genomic regions associated with the phenotype. To address LD, an ML model is trained on genotype and phenotype data, followed by analysis to identify the causal gene within the locus (Figure 1B). The architecture of the ML model allows direct identification of pleiotropic genes (Figure 1C). Incorporating transcriptomic data within a GSMM and GRN pipeline enables a mechanistic understanding of the GP relationship (Figure 1D). Our model is also tested on unseen data to better assess its predictive reliability (Figure 1E).

### Genotype encoding generation

We used three previously published yeast QTL mapping datasets^1,20,21^. All three datasets have been derived from a cross between BY and RM strains. BY (BY4741) is a standard laboratory strain initially derived from a soil isolate, and RM (RM-11a) is derived from a vine isolate with approximately 50,000 SNPs differing between one another. In this manuscript, we have named the above datasets as Bloom2013^20^, Bloom2015^21^, and Bloom2019^1^.

Genotype encoding was used to capture the genomic variation present in each yeast strain (Supplementary Figure 1A). We restricted the analysis to protein-coding regions of the genome, as most cellular functions are mediated by proteins and mutations in non-coding regions are more difficult to interpret functionally. Variant information was obtained from Variant Call Format (VCF) files and annotated using SnpEff^50^, which classifies variants based on their genomic position relative to the reference genome (R64-1-1.99) and predicts their functional consequences.

Synonymous variants, which do not alter the encoded protein sequence, were removed using SnpSift^51^. Each remaining variant was assigned an encoding score according to its predicted impact (Supplementary Table S2). Variant-level encodings were then aggregated at the gene level by assigning each gene the maximum encoding score among all its variants. This approach prioritizes mutations with higher predicted functional impact; for example, if a gene harbors both a non-synonymous mutation (encoding 1) and a nonsense mutation (encoding 2), the gene is assigned an encoding of 2, reflecting the greater likelihood that a nonsense mutation causes functional disruption via premature protein truncation. This procedure was repeated independently for each strain, yielding a gene-by-strain matrix of genotype encodings.

### Identifying QTLs

Identification of quantitative trait genes (QTGs) requires both genotype and phenotype information. For this analysis, the genotype data were represented as a binary vector for each gene, where 1 indicates the presence of at least one non-synonymous or loss-of-function mutation in that gene, and 0 indicates its absence. Phenotypes were binarized into high- and low-growth classes as described above.

For each gene, we constructed a contingency table relating mutation status to growth class (Supplementary Table S1) and applied Fisher’s exact test to assess whether mutation status was associated with the growth phenotype. To quantify the direction and magnitude of this association, we computed the odds ratio:

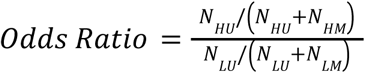

where *N*_*HU*_ and *N*_*HM*_ denote the numbers of high-growth strains without and with mutations, respectively, and *N*_*LU*_ and *N*_*LM*_ denote the corresponding counts for low-growth strains without and with mutations, respectively. An odds ratio greater than 1 indicates that mutations in the gene are associated with poorer growth, whereas an odds ratio less than 1 indicates an association with improved growth. *P* values from Fisher’s exact test were corrected for multiple hypothesis testing using the Benjamini–Hochberg procedure^52^.

### Chemical feature generation

Chemical features constituted the second input to the machine learning model, alongside genotype information. We generated these features using a previously described framework developed by our group^23^ (Supplementary Figures S1B and S2). For each molecule in the dataset, we first identified predefined functional groups (FGs), which correspond to chemically meaningful substructures with well-characterized properties (e.g., amines and carboxylic acids).

In addition to curated functional groups, we extracted mined functional groups (MFGs) directly from the SMILES representations of the molecules. MFGs correspond to frequently occurring substructures across the dataset and are not restricted to motifs with known chemical functions. The FG and MFG representations were concatenated and provided as input to an autoencoder neural network.

The autoencoder, which was pretrained on the PubChem database^53^, learns a compressed latent representation that captures salient structural and chemical information from the combined feature set. The resulting latent vectors were used as the chemical feature representation for downstream modeling.

### Phenotype processing

We formulated phenotype prediction as a binary classification problem, distinguishing strains that exhibit poor versus robust growth under a given chemical condition (Supplementary Figure S1C). Casting the task in this manner allows the approach to generalize across heterogeneous growth readouts, including yield or colony size measurements and sequencing-based bulk fitness assays.

To reduce ambiguity in class assignment, we restricted the analysis to strains displaying *extreme phenotypes*, defined as those lying in the tails of the phenotype distribution for each chemical condition. This strategy enables a more confident separation of high- and low-growth strains than would be possible with the full dataset.

For each condition, we computed the mean (μ) and standard deviation (σ) of the phenotype values. Strains were labeled according to a threshold parameter α as follows:

- strains with phenotype x < μ − ασ were assigned to the low-growth class (label 0);
- strains with phenotype x > μ + ασ were assigned to the high-growth class (label 1);
- strains with intermediate values were excluded from further analysis.

We evaluated α ∈ {0.25, 0.5, 1} and selected α = 0.5 because it provided a favorable trade-off between class separation and the number of retained training samples.

### Model Training

We tested four models to understand the GP relationship: Logistic Regression, Support Vector Machines (SVM), Random Forests, and Gradient Boosted Decision Trees (GBDT). The first two models were trained with the Scikit-Learn^54^ Python package, and the GBDTs and random forests were trained using the LightGBM package^55^. The models were selected across the range of complexities to test how informative our features are. We used the Optuna hyperparameter optimization framework^56^ to tune our models. We used the Tree Parzen Estimator (TPE) as the optimization algorithm. We ran the algorithm for 50 trials on the training data to determine the best hyperparameters. Details of all hyperparameters tuned across all models are provided in Supplementary Section 3.1.

Final model performance was evaluated using 5-fold cross-validation, with accuracy, area under the receiver operating characteristic curve (AUC–ROC), and F1 score as evaluation metrics. We observed that GBDTs performed best across the four models (Supplementary Figure S3A) and were selected for further downstream analysis.

### Interpretability Analysis

Once the models were trained, we performed interpretability of the best models using the SHAP algorithm from the SHAP Python package (version 0.45.1)^57^. SHAP values were used to quantify the contribution of individual features to model predictions, enabling the identification of genes with a strong influence on phenotype classification. Since the models were trained using 5-fold cross-validation, SHAP analysis was performed on each fold to obtain feature importance scores, which were averaged across folds to obtain robust estimates of gene importance. Genes highlighted by the interpretability analysis were subsequently analyzed using gene set enrichment analysis (GSEA)^58^ to gain functional insight.

### Gene set enrichment analysis

We performed GSEA to understand the functions of genes that were identified by the interpretability methods. GSEA aims to identify specific groups of genes (GO pathways, Cellular Components, KEGG pathways) that are enriched within a given set of genes. We used the enrichGO function from the clusterProfiler (version 4.10.1)^59^ and gProfiler2 (version 0.2.3)^60^ R packages to implement GSEA. We used a *p*-value cutoff of 0.05 with a Benjamini-Hochberg correction^52^ to identify significantly enriched groups within a given set of genes. We calculated the enrichment fold change to quantify enrichment.

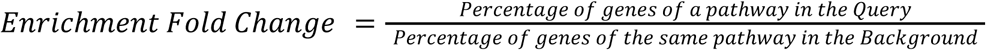

### Building strain-specific genome-scale metabolic models

Genome-scale metabolic models (GSMMs) are in silico representations of organisms’ metabolisms and are defined by the metabolic reactions that occur in the cell^41^.

To build strain-specific models, we started with the yeast-GEM (version 8.4.0)^38^, a consensus model of yeast metabolism and a transcriptomic dataset of the Bloom2013 strains grown in YNB media^39^. The model was built using MATLAB (version R2023a) and CobraToolBox (version 3.0)^61^. The model contained 2,806 metabolites, 4,131 reactions, and 1,063 genes. The yeast-GEM model has a glucose-limited minimal medium by default. We modified the media to closely match the Yeast Nitrogen Base (YNB) media from a previous study^39^ by setting the lower bound of the reactions to −10 (Supplementary Table S8). We could not fully represent the data because there were no exchange reactions involving iodide, ferric, or borate ions. We used three algorithms, INIT^62^, GIMME^63^, and SprintCore^64^, to build the strain-specific GSMMs. To obtain the most representative models, the Spearman correlation between the predicted fluxes through the biomass reactions and the experimental growth values was computed. The GIMME algorithm had the highest Spearman correlation of 0.31 (Supplementary Figure S9), and hence, the GSMMs built by the GIMME algorithm were selected for further analysis (For more details, see Supplementary Section 4).

We performed Flux Balance Analysis (FBA) to determine the optimal growth rates (flux through the biomass reaction) for each strain. Later, to binarize the strains’ phenotypes into high- or low-growth, we considered strains with growth rates below the mean as low-growing and those above the mean as high-growing. Following this, we compared the performance of the ML model and GSMM towards predicting the growth phenotype (Supplementary Table S9). The model analysis was restricted to strains accurately predicted by both the ML and GSM models (∼150 models).

### Gene-level analysis

We performed parsimonious flux balance analysis (pFBA)^40^ to identify which genes were responsible for the phenotype change. pFBA attempts to identify genes that are important for the growth of the organism and aims to find an FBA solution that minimizes the total flux through the model. In this sense, this gives the most efficient solution to the system. Following this, the model classifies the genes of the organism based on their effects on the system (Supplementary Table S3). pFBA was performed using the pFBA function from CobraToolBox^61^.

The identified genes were used as input for GSEA, which will inform us about the various subsystems enriched in the set.

### Reaction level analysis

The built strain-specific models identified reactions that were consistently present in high-growing strains and low-growing strains. To understand the distribution of the reaction fluxes of individual strains, flux sampling was performed using the OPTGP sampling algorithm^65^ from the CobraPy (version 0.29.0) package^66^.

Flux fold change for a reaction was calculated between the low- and high-growing phenotypes to understand the differences in reaction rates between the two conditions.

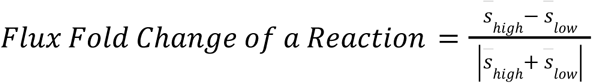

where *S̅* refers to the mean flux through the particular reaction. The cutoff of 0.82 was chosen to identify the reactions. The metric and cut-off were chosen from a previous study^41^. If the fold change is greater than 0.82, they are grouped under reactions up-regulated in high-growing strains and vice versa. Following this, we performed an enrichment analysis for the reactions to identify if any pathways were highly significantly overrepresented.

### Gene regulatory network analysis

We used the BioNERO (version 1.10.3) R package^43^ to build and analyze the gene regulatory network (GRN) from the data^39^. The TPM data were preprocessed by removing low-expression genes (all genes with average expression <10 TPM) and low-variance genes (variance < the 10th percentile of all gene variances). The GRN was constructed using the *exp2grn* function. The transcription factor information used to build the GRN was obtained from the YEASTRACT database^42^. The edges obtained were filtered using the *grn_filter* function to ensure that the resultant graph follows a scale-free topology.

## Supporting information

Supplementary File 2- pFBA Enrichment High Genes

Supplementary File 1 - Bloom2013 QTL Analysis

## DATA AND CODE AVAILABILITY

The data files used for all analyses, from model training to GSM and GRN analysis, along with the Supplementary Information and tables, are available on Zenodo (link). Codes for reproducing all the results shown in the paper are available on GitHub https://github.com/bisect-group/yeast_growth_analysis.

## ACKNOWLEDGEMENTS

We thank members of IBSE and WSAI for their valuable inputs and suggestions. We thank members of the BiSeCT and Systems Genetics Labs for their inputs.

## AUTHOR CONTRIBUTIONS

NB and HS conceptualized the project; RMRL performed all analysis and modelling and wrote the first draft; RMSB developed tools for the machine learning analysis; NB and HS supervised the project and acquired funding; and all authors revised, edited, and finalized the manuscript.

## SUPPLEMENTARY INFORMATION

**Supplementary Information** contains Supplementary Tables S1-S9 and Supplementary Figure S1-S8.

**Supplementary File 1**: SHAP resolved QTGs from every QTL previously identified in the Bloom2013 dataset^20^.

**Supplementary File 2**: The pathway enrichment result for all pFBA genes unique to the high-growing strains.

