## Supplementary Information for "Interpretable machine learning meets systems biology to decode genotype-phenotype maps"

### 1 Supplementary Tables

Table S1: Contingency Table of gene mutation and growth phenotype

| **Growth** | **Unmutated** | **Mutated** |
|---|---|---|
| High | $N_{HU}$ | $N_{HM}$ |
| Low | $N_{LU}$ | $N_{LM}$ |

Table S2: The Possible Gene-level Encodings

| **Type of Mutation** | **Encoding** |
|---|---|
| No mutation or Synonymous | 0 |
| Missense | 1 |
| Nonsense | 2 |
| Frameshift | 3 |

Table S3: Gene Classification of pFBA

| **Category** | **Description** |
|---|---|
| Essential | Growth not possible without these |
| Optimal | Genes whose reactions carry a nonzero flux at optimal growth |
| Metabolically Less Efficient (MLE) | Genes whose reactions reduce the growth of the organism |
| Enzymatically Less Efficient (ELE) | Genes whose reactions increase the total flux through the system |

#### 2 Supplementary Figures

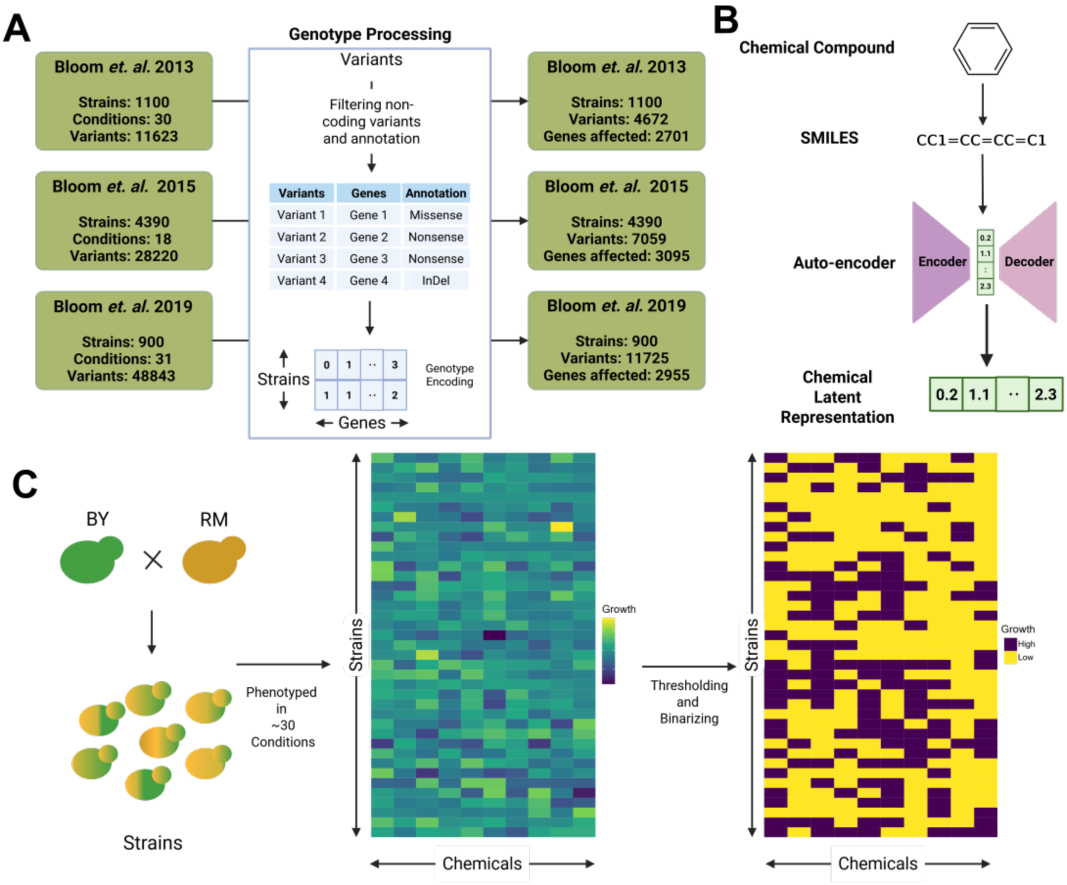

Figure S1: (A) We obtained segregant data from the BY and RM cross from three publications. Condition here refers to the different chemicals the strains were grown in. We applied our genotype encoding generation framework on the variant data from these datasets. (B) The deep learning pipeline to represent our chemicals in a numerical form. (C) Binarizing growth phenotype to get qualitative measurements of growth.

#### 3 Machine learning models integrating genotype and chemical information

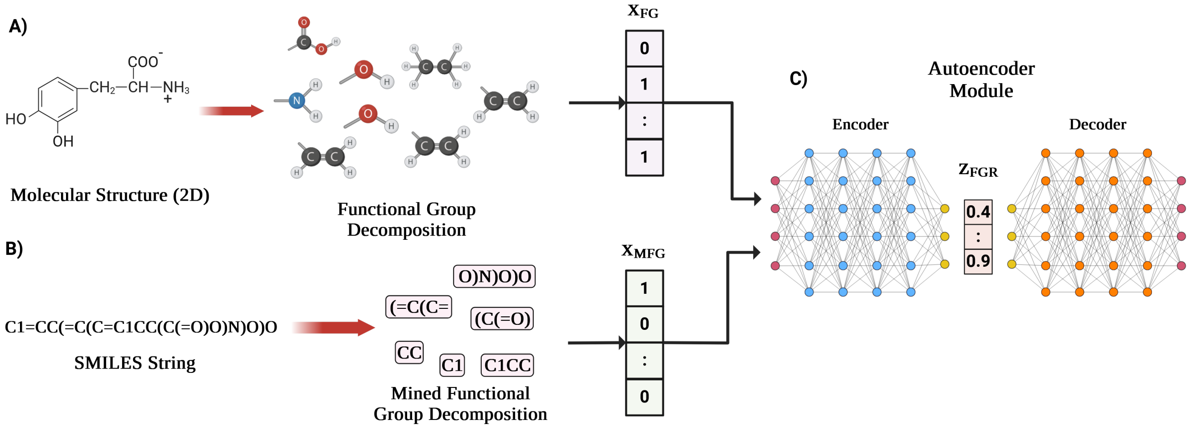

Figure S2: The pipeline used to generate latent representation of the chemical molecules. (A) From the structure of the molecule, we extract the functional groups that are present in the molecule. This results in a binary vector $\mathbf{X}_{FG}$ which encodes the presence or absence of a given functional group. (B) From the SMILES string of the given molecule, we extract the mined functional groups, which are commonly occurring sub-strings in SMILES representation. This will yield us another binary vector $\mathbf{X}_{MFG}$ (C) We concatenate the $\mathbf{X}_{FG}$ and $\mathbf{X}_{MFG}$ as pass it as the input to an autoencoder neural network. The encoder produces a latent representation $\mathbf{Z}_{FGR}$ which will be used as the input for the downstream training.

#### 3.1 ML model Hyperparameters for Tuning

##### 3.1.1 Logistic Regression

Table S4: Logistic Regression Hyperparameters

| Name | Description | Values/Range[a] |
|---|---|---|
| `penalty` | Type of penalization to be applied | {`l1, l2, elasticnet`} |
| `l1_ratio` | Ratio of l1 regularization to l2 [b] | `[0,1]` |
| C | Penalization strength | `[1e-3,1e3]` |

[a] [] represents range, {} represent all possible values
[b] Active only if `penalty=elasticnet`

##### 3.1.2 Support Vector Machines

Table S5: SVM Hyperparameters

| Name | Description | Values/Range[a] |
|---|---|---|
| `C` | Regularization strength | `[1e-3,1e3]` |
| `kernel` | Type of kernel to use | {`linear, poly, rbf, sigmoid`} |
| `gamma`[b] | Coefficient for `rbf, poly` and `sigmoid` kernels | {`scale, auto`} |
| `degree`[c] | Degree | `[2,7]` |

[a] [] represents range, {} represent all possible values
[b] Coefficients for `rbf`, `poly`, and `sigmoid` kernels
[c] Only active if `kernel=poly`

##### 3.1.3 Random Forest

Table S6: Random Forest Hyperparameters

| Name | Description | Values/Range[a] |
|---|---|---|
| `n_estimators` | Number of trees to be built | `[10,1000]` |
| `split_criterion` | Loss function | {`gini, entropy`} |
| `max_samples` | Fraction of samples to be considered for tree building | `[0.1,1]` |
| `max_features` | Fraction of features considered for tree building | `[0.1,1]` |
| `max_depth` | Maximum tree depth | `[2,12]` |
| `min_samples_leaf` | Minimum number of samples at a leaf node | `[5,100]` |

[a] [] represents range, {} represent all possible values

##### 3.1.4 Gradient-Boosted Trees

Table S7: Gradient-Boosted Trees Hyperparameters

| Name | Description | Values/Range[a] |
|---|---|---|
| `n_estimators` | Number of trees to be built | `[10,1000]` |
| `lambda_l1` | L1 penalty | `[0,1]` |

| | | |
|---|---|---|
| lambda_l2 | L2 penalty | [0,1] |
| num_leaves | Total number of leaves in the model | [2,256] |
| feature_fraction | Fraction of features to be considered for tree building | [0.1,1] |
| bagging_fraction | Fraction of samples to be considered for tree building | [0.1,1] |
| bagging_freq | Frequency for bagging | [2,7] |
| min_child_samples | Minimum number of samples at a leaf node | [5,100] |
| learning_rate | Learning rate for boosting | [1e-3, 1e-1] |

[a][] represents range, {} represent all possible values

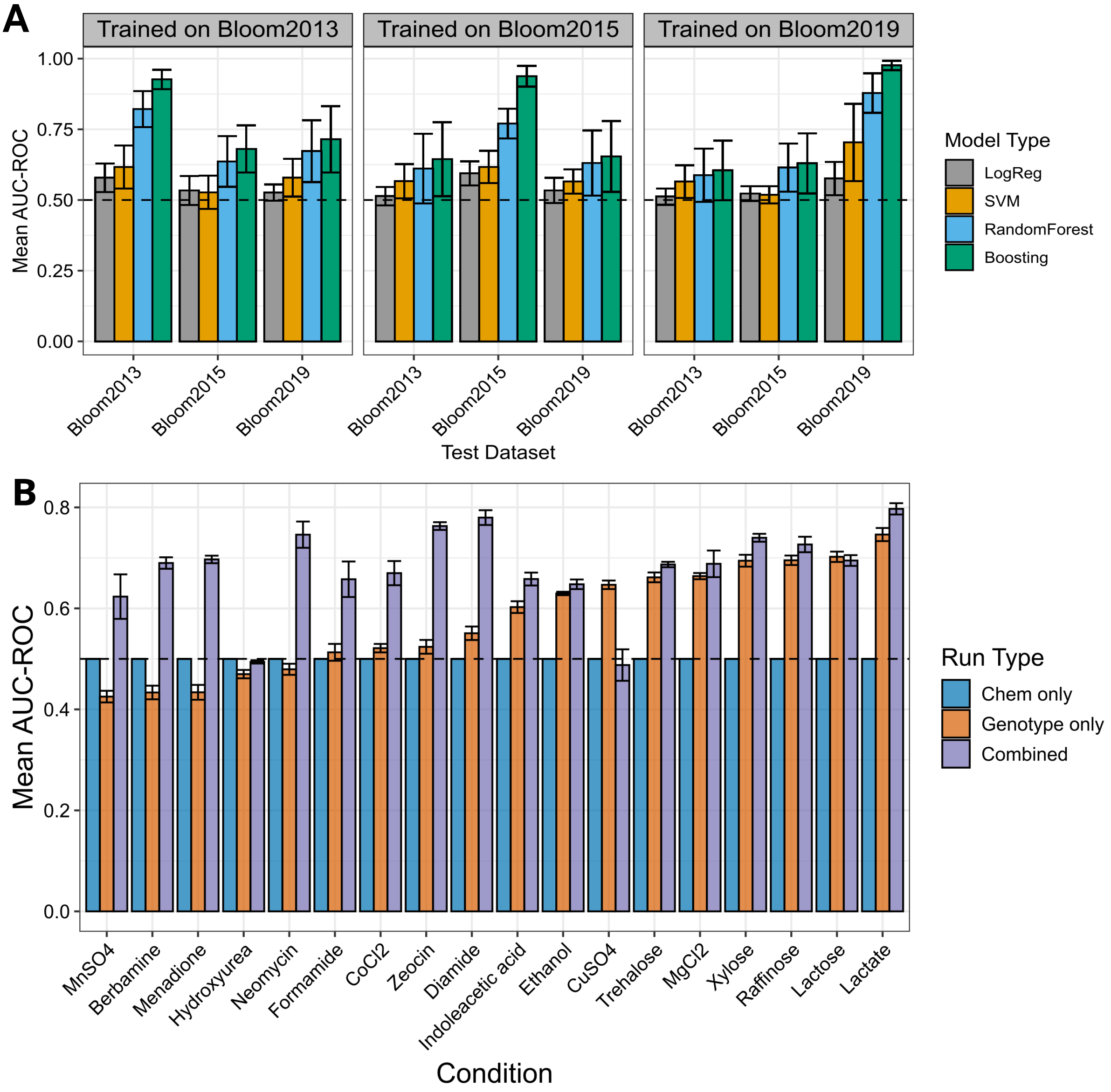

Figure S3: (A) Performance of the models used in this study compared to a random model measured using AUC-ROC metric. Each point on the horizontal axis represents an experiment where the model was trained on a particular dataset and tested on a different dataset. Except for the logistic regression model, all the models perform much better than the random model when it comes to predicting the growth. (B) The condition-wise performance of Gradient-Boosted Decision Tree Model trained on Bloom2013 data and tested on Bloom2015 dataset. We observed a variation in the performance of both the combined and genotype-only models. While in some carbon sources, the genotype model is able to perform just as well as the combined model, it struggles to perform in diverse chemical conditions.

#### 3.2 Performance comparisons - all combinations

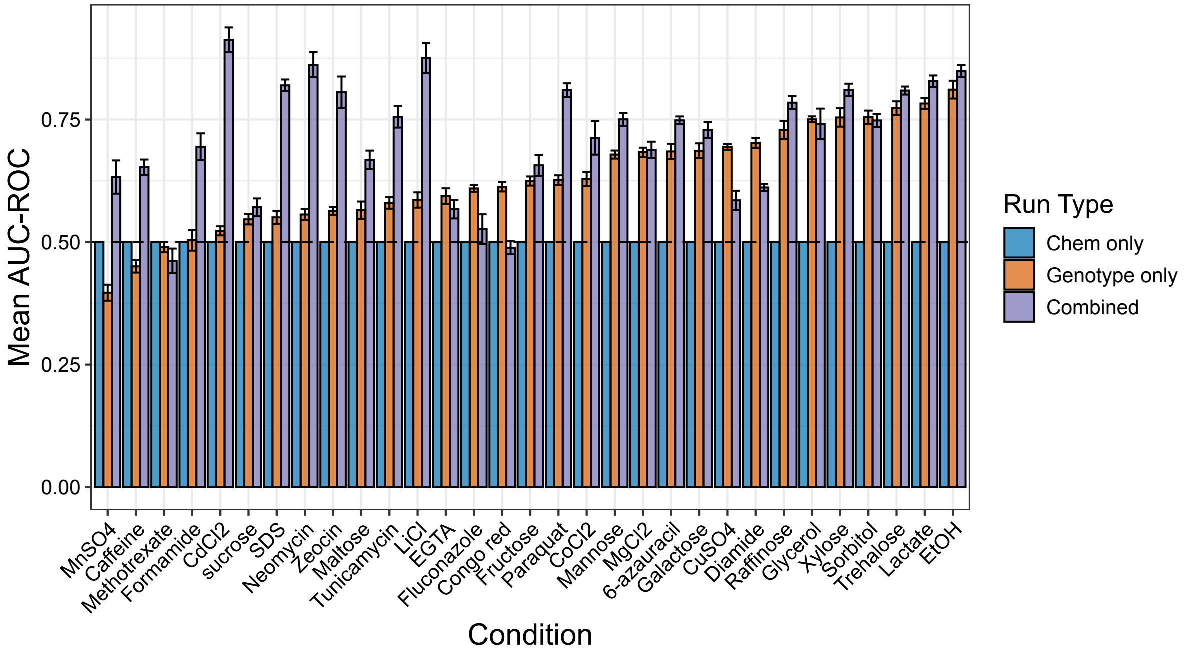

Figure S4: Condition-wise performance of Bloom2013 trained model on Bloom2019 dataset

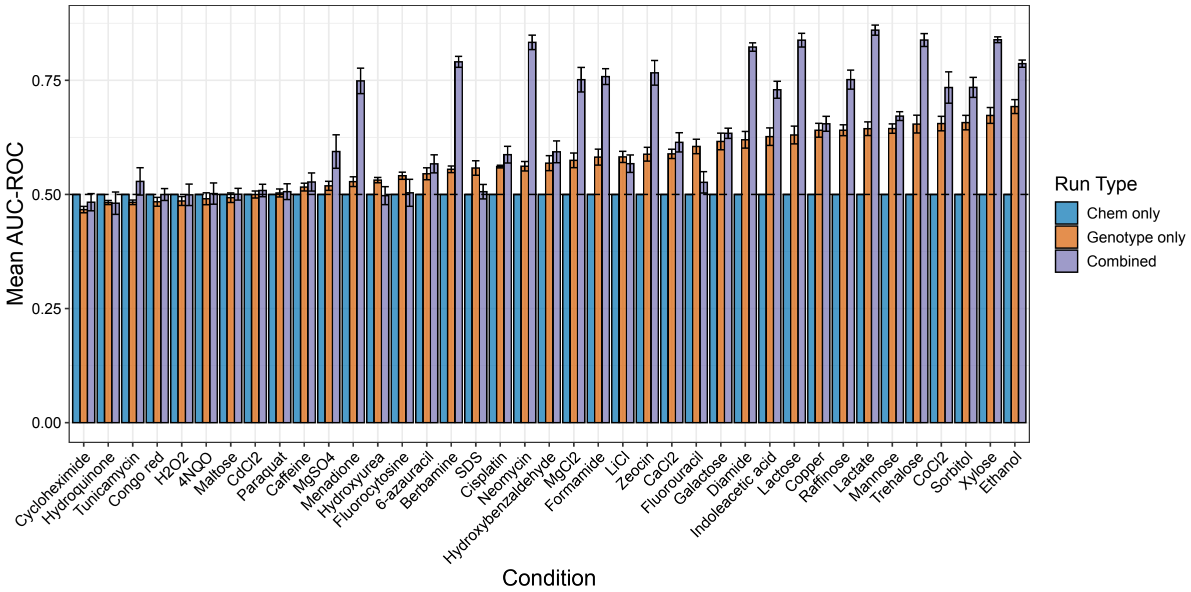

Figure S5: Condition-wise performance of Bloom2015 trained model on Bloom2013 dataset

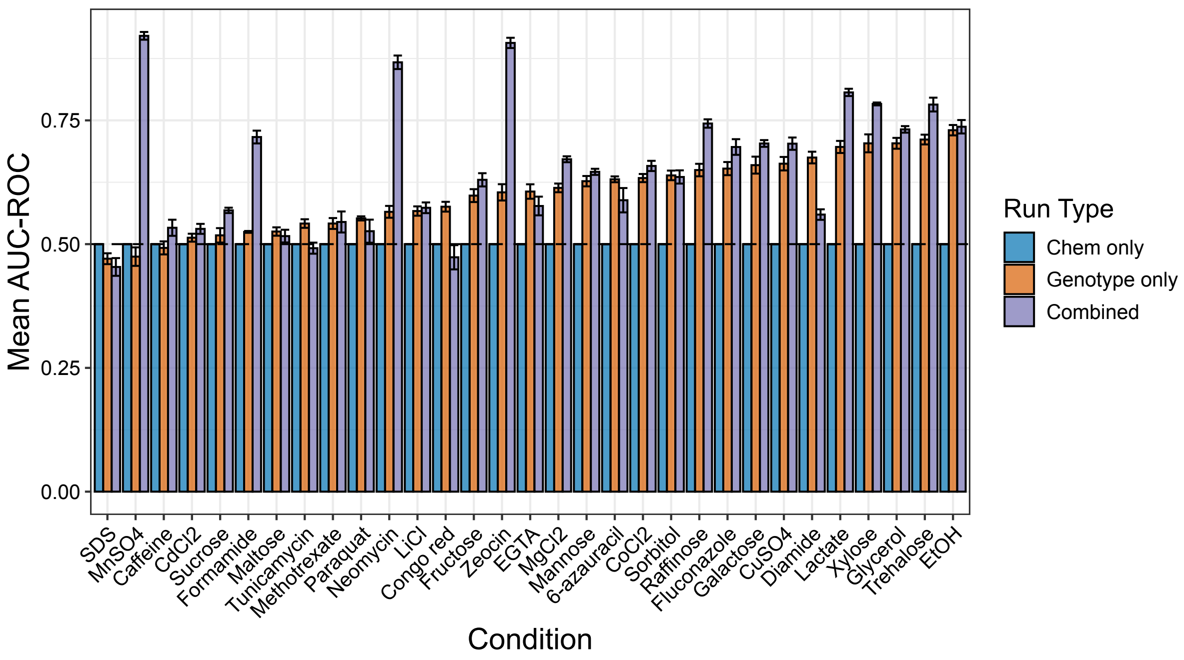

Figure S6: Condition-wise performance of Bloom2015 trained model on Bloom2019 dataset

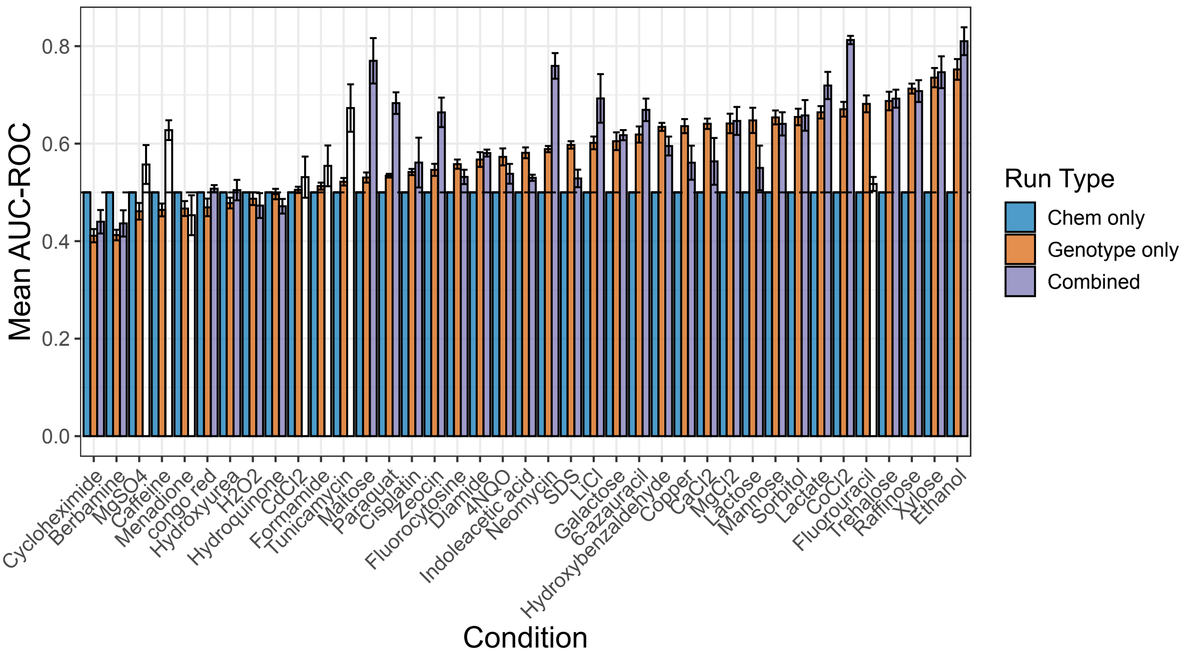

Figure S7: Condition-wise performance of Bloom2019 trained model on Bloom2013 dataset

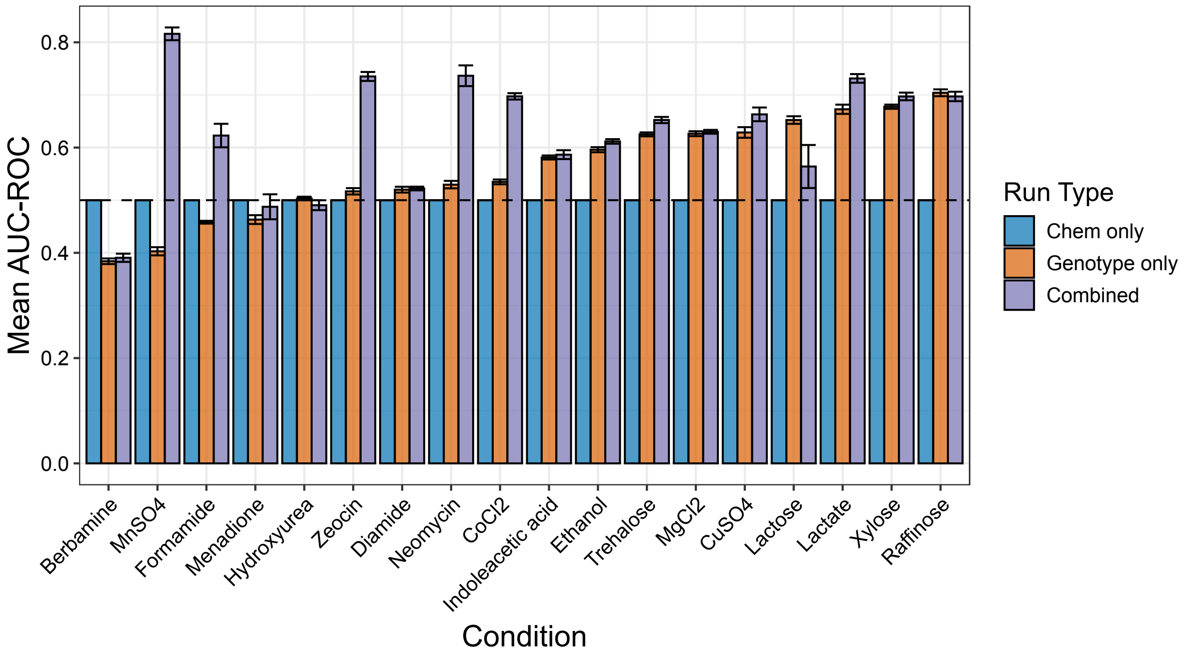

Figure S8: Condition-wise performance of Bloom2019 trained model on Bloom2015 dataset

### 4 GSMM Modelling

#### 4.1 Building and validation

We built strain-specific GSMMs for Bloom2013 strains using data from Albert et al., 2018. Prior to model building, the GSMM exchange reaction was modified to best represent the growth media in Albert et al., 2018. The details of the exchange reactions are given in Table S8. Strain-specific model building involves identifying the active genes and reactions for a given context, followed by modifying the GSMM to reflect the same. The transcriptomic data from Albert et al., 2018 are specific to each strain in Bloom2013 and we used this data as a starting point. The process of identifying active genes is called thresholding. The thresholding algorithm that we used was LocalGini (Pavan Kumar & Bhatt, 2023). To identify the active reactions, we utilized the Gene-Protein-Reaction (GPR) rules of the GSMM, which are a set of boolean rules that govern which reactions will be active given a set of genes. It is possible that the models described only by the active reactions are incomplete (reactions without a GPR rule may not be considered), therefore, a model-building algorithm will add the minimal number of reactions to make the model complete. There are several model-building algorithms that can build the final context-specific model. Three model extraction algorithms were chosen for experimentation: GIMME(Becker & Palsson, 2008), SprintCore (**SprintFamily**) and INIT (Agren et al., 2012). Of the three model extraction methods, we had chosen, INIT failed to provide models that grow even though the biomass reaction was provided as a core reaction. Therefore the rest of the analysis was limited to GIMME and SprintCore built models.

Table S8: The components of YNB media and the reactions modified to represent it for the Yeast-GEM model

| Component | Reaction ID | Reaction Name |
|---|---|---|
| **Carbon source** | | |
| Glucose | r_1714 | D-glucose Exchange |
| **Vitamins** | | |
| Biotin | r_1141 | Biotin Exchange |
| Pantothenate | r_1548 | R-pantothenate Exchange |
| Folate | r_1792 | Folic acid Exchange |
| Inositol | r_1947 | Myo-inositol Exchange |
| Nicotinic acid | r_1967 | Nicotinate Exchange |
| p-Aminobenzoic acid | r_1604 | 4-Aminobenzoate Exchange |
| Pyridoxine | r_2028 | Pyridoxine Exchange |
| Riboflavin | r_2038 | Riboflavin Exchange |
| Thiamine | r_2067 | Thiamine (1+) Exchange |
| **Salts** | | |
| Copper (2+) | r_4594 | Cu (2+) Exchange |
| Manganese (2+) | r_4595 | Mn (2+) Exchange |
| Sodium | r_2049 | Sodium Exchange |
| Potasium | r_2020 | Potassium Exchange |
| Phosphate | r_2005 | Phosphate Exchange |
| Zinc (2+) | r_4596 | Zn (2+) Exchange |
| Magnesium (2+) | r_4597 | Mg (2+) Exchange |
| Calcium (2+) | r_4600 | Ca (2+) Exchange |
| Ammonium | r_1654 | Ammonium Exchange |
| Sulphate | r_2060 | Sulphate Exchange |

| | | |
|---|---|---|
| Chloride | `r_4593` | Chloride Exchange |

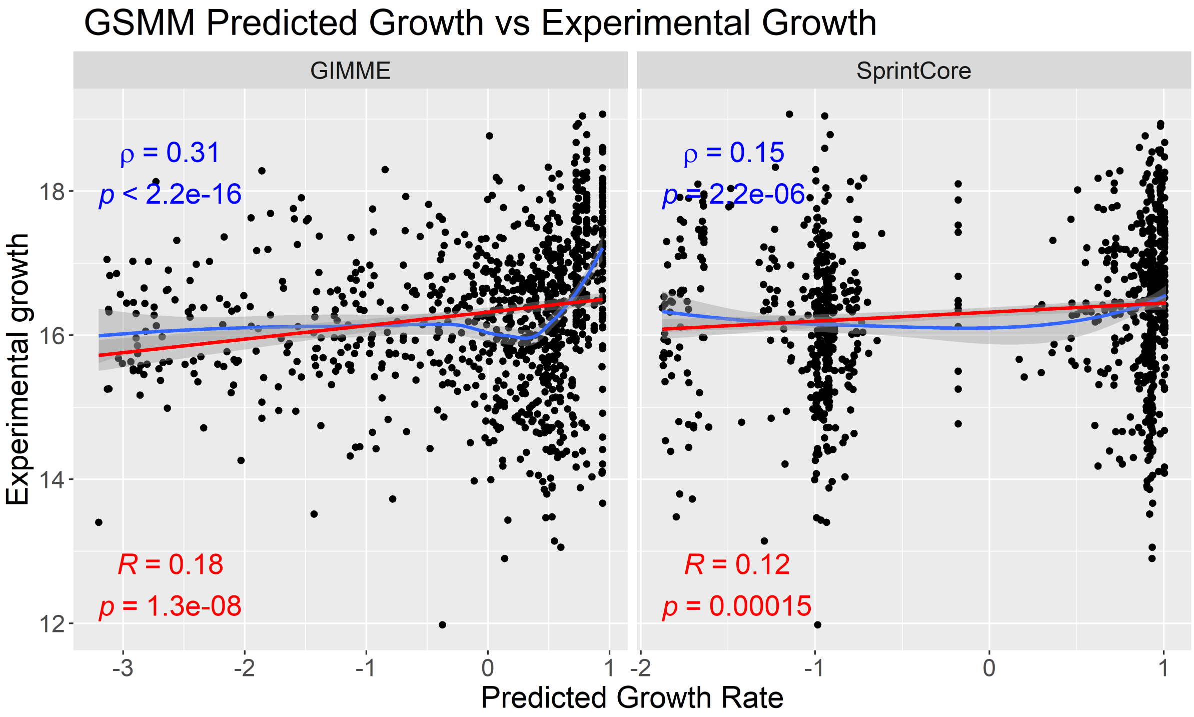

Figure S9: We compared models built by SprintCore and GIMME algorithms to identify the most representative model. On the X-axis is the predicted growth rate which was the flux through the biomass reaction. On the Y-axis is the colony size for each strain. An ideal model will have its predicted growth rate correlate highly to the experimental values. The values marked in red is the Pearson correlation between the data in the two axes and the blue value is the Spearman correlation.

Table S9: Comparing the concordance between the different datasets. ML refers to an Gradient-Boosted Decision Tree trained on the Bloom2013 dataset. GSMM refers to the binarized growth rate from Genome Scale Metabolic Models.

| Data 1 | Data 2 | Concordance |
|---|---|---|
| Experimental | ML prediction | 70.67% |
| Experimental | GSMM SprintCore | 58.67% |
| Experimental | GSMM GIMME | 70.67% |
| GSMM SprintCore | ML prediction | 50.67% |
| GSMM GIMME | ML prediction | 61.33% |
| GSMM GIMME | GSMM SprintCore | 56.00% |

After binarizing the predicted growth rate of the strains, we looked at the concordance between GSMM prediction and the ML model prediction and observed that GIMME performed better than SprintCore-built models (Table S9). We therefore used only the GIMME based models for further analysis.
